# Characterization of a lethal Crimean-Congo hemorrhagic fever virus infection model in interferon-deficient mice for studies of virus pathogenicity and vaccine’s efficacy

**DOI:** 10.64898/2025.12.23.696184

**Authors:** Andrey E. Siniavin, Anna A. Iliukhina, Daria M. Grousova, Ilya D. Zorkov, Inna V. Dolzhikova, Vladimir A. Gushchin

**Affiliations:** Federal State Budget Institution “National Research Centre for Epidemiology and Microbiology Named af-ter Honorary Academician N F Gamaleya” of the Ministry of Health of the Russian Federation, 123098 Moscow, Russia; Federal State Autonomous Educational Institution of Higher Education “I.M. Sechenov First Moscow State Medical University” of the Ministry of Health of the Russian Federation (Sechenov University), 119991 Moscow, Russia; Department of Virology, Lomonosov Moscow State University, 119991 Moscow, Russia

## Abstract

Crimean-Congo hemorrhagic fever virus (CCHFV) is a highly pathogenic tick-borne virus causing severe human disease and representing a significant public health threat. The absence of licensed vaccines and specific antiviral therapies highlights the need for well-characterized experimental models suitable for studying viral pathogenicity and evaluating medical countermeasures. In this study, we established and characterized an integrated in vitro and in vivo platform for CCHFV research. Virus propagation was optimized by comparing Vero E6 and SW-13 cell lines. SW-13 cells supported efficient CCHFV replication resulting in pronounced virus-induced cytopathic effects, whereas no detectable cytopathic effects were observed in Vero E6 cells. Virus titers determined by cytopathic effect based endpoint dilution and antigen ELISA coincided, supporting the use of SW-13 cells for virus stock preparation. Using virus stocks generated in SW-13 cells, interferon-deficient mice were infected with different doses of CCHFV to assess dose-dependent pathogenicity. Infection resulted in severe disease and high mortality across all tested doses. Based on survival analysis, a dose of 10 TCID50 was selected for further characterization, which revealed progressive weight loss and high viral loads in peripheral organs. Overall, this study establishes a robust IFN knockout mouse model suitable for studying CCHFV pathogenicity, comparing viral genovariants, and evaluating vaccines, antiviral compounds, and neutralizing antibodies.

## Introduction

Crimean-Congo hemorrhagic fever (CCHF) is a severe tick-borne zoonotic disease caused by Crimean-Congo hemorrhagic fever virus (CCHFV), a member of the genus *Orthonairovirus* within the family *Nairoviridae* [1]. Human infection is characterized by acute febrile illness, systemic inflammation, hemorrhagic manifestations, and high case fatality rates that may reach 30-40% in endemic regions [2]. CCHFV is widely distributed across Africa, the Middle East, Eastern and Southern Europe, and Asia, and its geographical range continues to expand, likely driven by climate change, vector distribution, and increased contact between human and animal [3]. These factors underscore the urgent need for effective vaccines and antiviral countermeasures against CCHFV.

CCHFV exhibits remarkable genetic diversity, with multiple phylogenetically distinct genotypes circulating in different geographic regions [4]. Increasing evidence suggests that viral genetic variability may influence virulence, replication efficiency, immune evasion, and disease severity [4,5]. However, systematic comparative studies addressing the pathogenic potential of different CCHFV genovariants remain limited, largely due to the lack of robust and reproducible small animal models that support productive infection and lethal disease.

A critical feature of CCHFV pathogenesis is its ability to antagonize the host innate immune response, particularly type I interferon (IFN-I) signaling pathways [6]. In immunocompetent mice, intact interferon responses effectively restrict viral replication, resulting in subclinical or abortive infection [6]. Consequently, conventional mouse models are poorly suited for evaluating CCHFV virulence, vaccine efficacy, or antiviral activity. In contrast, mice deficient in type I interferon signaling, including IFN-I receptor knockout (IFNAR^−^/^−^) or STAT1-deficient mice, are highly susceptible to CCHFV infection and develop a systemic, often fatal disease that recapitulates key features of human CCHF, such as high viral loads in peripheral organs and rapid disease progression [6,7]. Interferon-deficient mouse models have therefore emerged as essential tools for CCHFV research and are increasingly used for preclinical evaluation of vaccine candidates and antiviral compounds [7,8].

Nevertheless, detailed dose–response characterization, survival kinetics, and organ-specific viral replication profiles remain insufficiently defined for many viral strains and genotypes. Such baseline characterization is particularly important when comparing genetically distinct CCHFV isolates or assessing the protective efficacy of vaccines against heterologous genovariants.

In the present study, we comprehensively characterize a lethal CCHFV infection model in IFN knockout mice by challenging animals with different viral doses and monitoring survival outcomes and viral burden in key target organs. This model provides a robust and reproducible platform for future studies aimed at dissecting the pathogenic determinants of distinct CCHFV genovariants, evaluating vaccine-induced protection, and assessing the efficacy of antiviral compounds. Ultimately, this approach will facilitate a better understanding of CCHFV virulence mechanisms and support the development of effective medical countermeasures against this high-consequence pathogen.

## Methods

### Cells and virus

Vero E6 (ATCC, CRL-1586) and SW-13 (ATCC, CCL-105) cells were maintained in Dulbecco’s modified Eagle’s medium (DMEM; Gibco, USA) supplemented with 10% fetal bovine serum (FBS; HyClone, USA) at 37°C in a humidified atmosphere containing 5% CO_2_.

The CCHFV strain 6/H-73 used in this study was obtained from the State Collection of Viruses (Gamaleya Center, Moscow, Russia). The virus stock was prepared from the supernatant of infected SW-13 cells under biosafety level 3 (BSL-3) conditions.

Viral RNA copy numbers and the 50% tissue culture infectious dose (TCID_50_) per milliliter of the virus stock were determined by quantitative reverse transcription polymerase chain reaction (RT-qPCR) and endpoint dilution titration, respectively. In addition, virus titers in infected cell culture supernatants were determined by antigen-based enzyme-linked immunosorbent assay (ELISA; VectoCrimea-KGL-antigen, Vector-Best, Russia).

### Mouse infection

Ifnar1^−^/^−^ (Ifnar1 KO, catalog number C001268) mice on a C57BL/6NCya genetic background were purchased from Cyagen (China). Animals were randomly assigned to experimental groups and were 6–8 weeks of age at the time of infection. Mice were infected intraperitoneally with different doses of CCHFV. Body weight and survival were monitored daily for 14 days post infection. Animals were humanely euthanized after reaching predefined humane endpoint criteria, including ataxia, severe lethargy (lack of response to touch), hemorrhagic discharge, tachypnea, limb paralysis, or body weight loss exceeding 30%.

### Viral Load Analysis

For the analysis of viral loads in organs, mice were humanely euthanized on day 3 post infection, and tissues were collected into homogenization tubes. Tissue samples were homogenized at 30 Hz for 1 min using a TissueLyser (Qiagen) and briefly centrifuged at maximum speed to pellet debris. Clarified homogenates were subsequently used for RNA extraction and quantitative RT-PCR analysis.

## Results

### Optimization of virus propagation for in vivo experiments

Prior to *in vivo* studies, preliminary experiments were conducted to optimize CCHFV propagation and virus stock preparation. Two cell lines commonly used for CCHFV replication, Vero E6 and SW-13, were infected and compared for their ability to support productive infection. CCHFV infection of SW-13 cells resulted in a pronounced virus-induced cytopathic effect (CPE), which became evident during the course of infection. In contrast, no detectable CPE was observed in infected Vero E6 cells under the same experimental conditions, despite successful infection. These observations indicated marked differences in virus-cell interactions between the two cell lines.

Virus titers obtained from infected SW-13 cells were determined using two independent methods: CPE-based endpoint titration (TCID_50_) and antigen detection by enzyme-linked immunosorbent assay (ELISA). Both approaches yielded comparable virus titers, demonstrating consistency between CPE-based and antigen-based quantification methods and confirming efficient virus accumulation in SW-13 cells.

Based on the robust cytopathic effect and reproducible high virus titers, SW-13 cells were selected as the optimal in vitro system for CCHFV propagation. This model was subsequently used for the generation of virus stocks for in vivo experiments and provides a suitable platform for future evaluation of antiviral compounds and neutralizing antibodies.

### Dose-dependent lethality of CCHFV in IFN knockout mice

To assess the dose-dependent pathogenicity of CCHFV, IFN knockout mice were infected intraperitoneally with 10, 100, or 1000 TCID_50_ of CCHFV strain 6/H-73, and survival was monitored for 14 days post infection (dpi). Infection with all tested doses resulted in lethal disease (Figure 1). Mice infected with the highest dose (1000 TCID_50_) exhibited the most rapid disease progression, with 100% mortality observed by day 5 post infection. Animals challenged with 100 or 10 TCID_50_ showed a slightly delayed onset of mortality; however, survival outcomes were comparable between these groups, with approximately 20% of animals surviving until the end of the observation period. These data indicate that CCHFV causes severe and largely dose-independent lethal disease in IFN-deficient mice across a broad range of inoculation doses.

**Figure 1.**
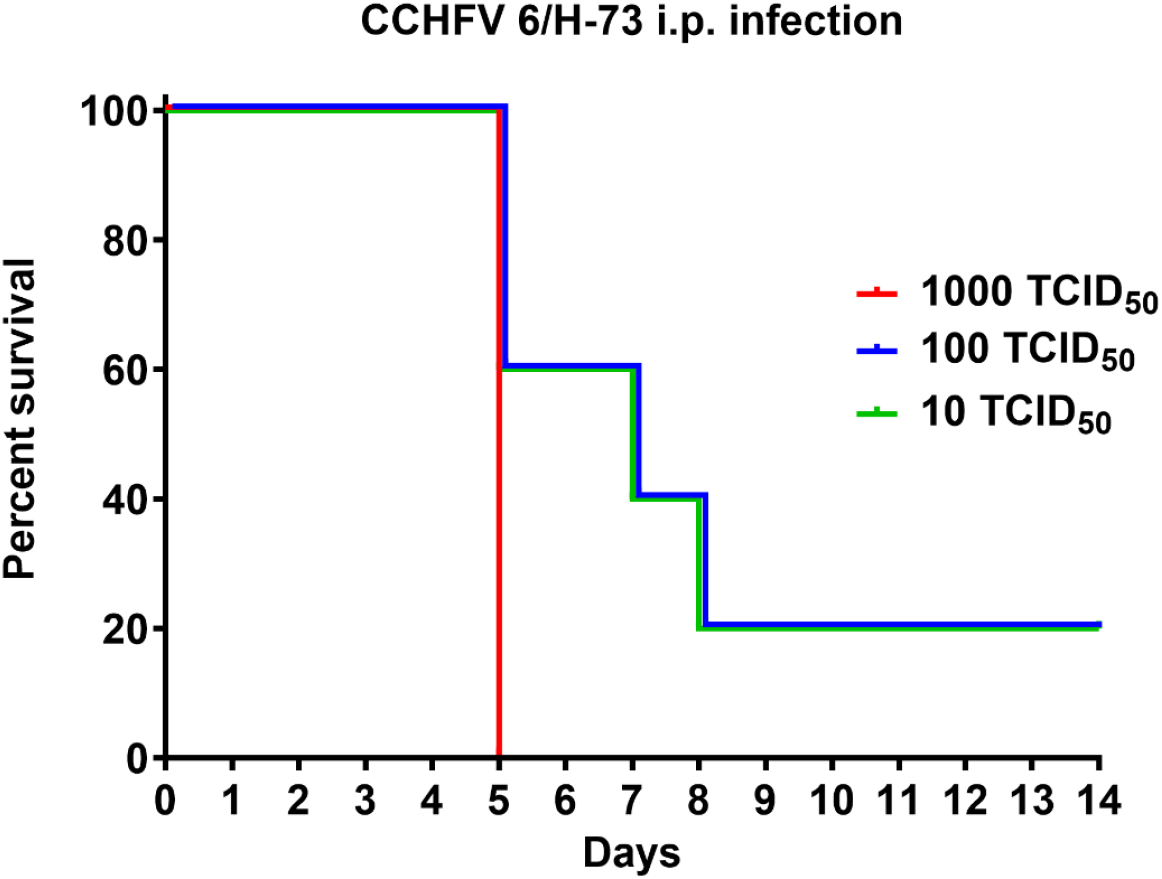
Dose-dependent survival of IFN knockout mice infected with CCHFV. Interferon knockout mice (n=5) were infected intraperitoneally with 10, 100, or 1000 TCID_50_ of CCHFV. Survival was monitored daily for 14 days post infection. Survival curves are shown as Kaplan– Meier plots.

### Pathological characterization of CCHFV infection in IFN KO mice after infectious dose optimization

Based on the survival analysis, a dose of 10 TCID_50_ was selected for further characterization of disease progression. Following infection, mice exhibited progressive and pronounced weight loss starting at day 2–3 post infection (Figure 2 A). Body weight steadily declined throughout the course of disease, reaching approximately 65–70% of the initial body weight by days 6–7 post infection, consistent with the onset of severe clinical disease. Weight loss preceded mortality and served as a reliable indicator of disease progression in this model (Figure 2 B).

**Figure 2.**
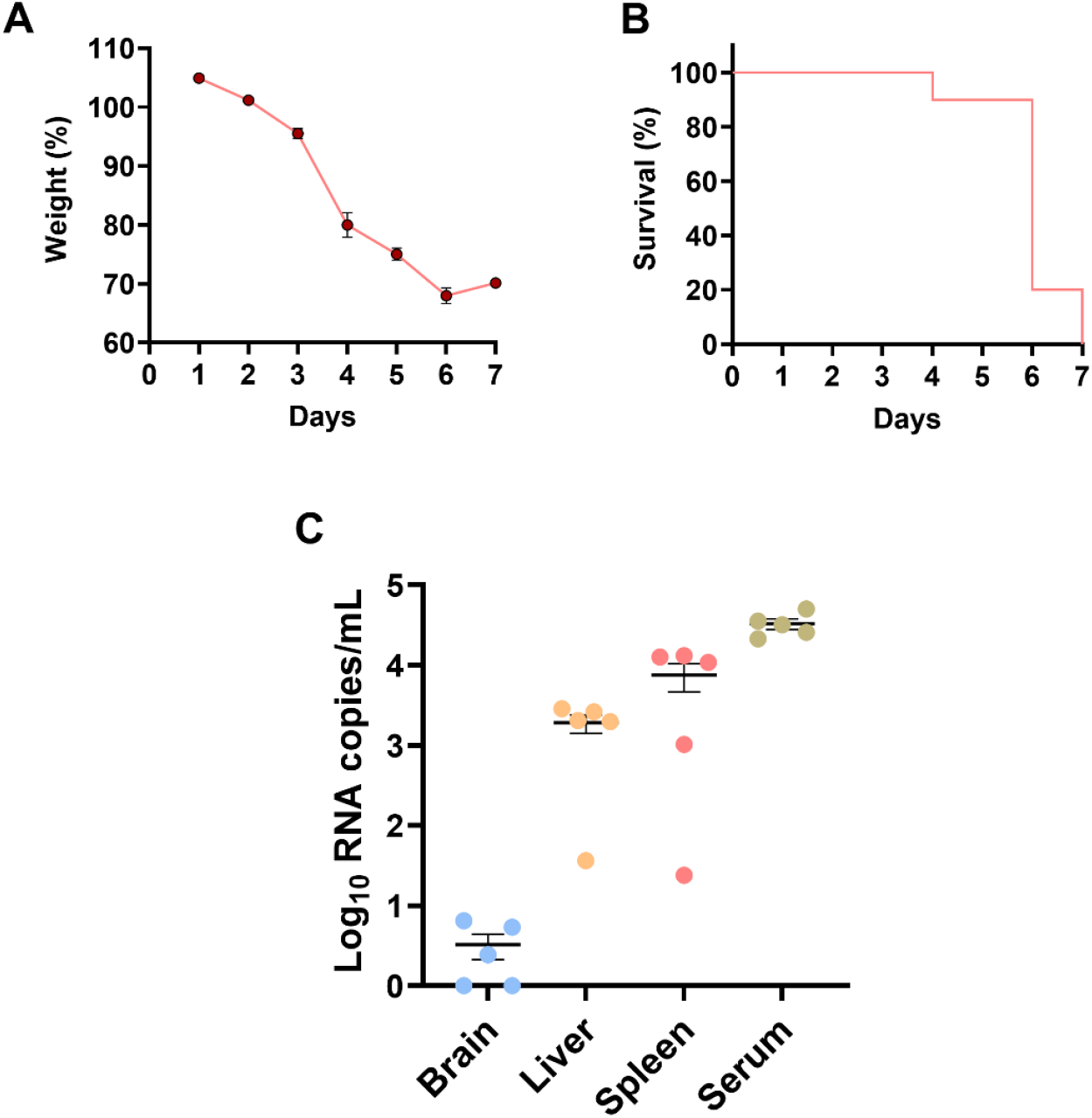
Disease progression and viral burden in IFN KO mice infected with CCHFV at the selected dose. IFN knockout mice were infected with 10 TCID_50_ of CCHFV. Body weight (A) and survival (B) was monitored daily. Data are presented as percentage of initial body weight. Viral RNA levels (C) were quantified in serum and selected organs of IFN KO mice infected with 10 TCID_50_ of CCHFV. The dots represent individual viral load values. Data are presented as mean with SEM.

To determine viral dissemination and tissue tropism, viral RNA levels were quantified in selected organs and serum of mice infected with 10 TCID_50_ of CCHFV. High viral loads were detected in peripheral organs, with the highest levels observed in serum and spleen, reaching approximately 4-5 log_10_ RNA copies/ mL (Figure 2 C). Substantial viral RNA levels were also detected in the liver, whereas viral loads in the brain remained comparatively low. These results demonstrate efficient systemic replication of CCHFV and widespread viral dissemination in IFN knockout mice following low-dose infection.

## Discussion

In the present study, we established and comprehensively characterized an experimental framework combining optimized *in vitro* propagation of CCHFV with a lethal IFN-deficient mouse model. This integrated approach provides a robust platform for studying CCHFV pathogenicity, genetic variability, and the efficacy of medical countermeasures, including vaccines, antiviral compounds, and neutralizing antibodies.

Efficient virus propagation is essential for reproducibility of *in vivo* studies. Our comparative analysis of Vero E6 and SW-13 cells demonstrated that SW-13 cells represent a superior *in vitro* system for CCHFV infection, supporting productive viral replication with a pronounced virus-induced cytopathic effect. In contrast, infection of Vero E6 cells did not result in detectable CPE under the same conditions, despite their widespread use for arbovirus propagation. These findings are consistent with previous reports identifying SW-13 cells as highly permissive for CCHFV replication and suitable for virus isolation and titration [7,9]. Importantly, virus titers determined by CPE-based endpoint dilution and antigen ELISA were similar, confirming the reliability of both quantification methods and supporting the use of SW-13 cells for standardized virus stock preparation, antiviral screening, and neutralization assays.

Using virus stocks generated in this optimized system, we characterized CCHFV infection in IFN knockout mice across a range of inoculation doses. Infection resulted in rapid and severe disease with high mortality at all tested doses, indicating that CCHFV pathogenicity in this model is largely dose independent. Similar lethal outcomes have been described in IFNAR^−^/^−^ and STAT1^−^/^−^ mice, highlighting the central role of type I interferon signaling in controlling CCHFV replication and preventing systemic disease [6,7,11,15]. The extreme susceptibility observed even at low infectious doses underscores the efficiency with which CCHFV exploits impaired innate immune responses to establish disseminated infection.

Progressive and pronounced weight loss was observed following low-dose infection and closely correlated with disease severity and mortality. Weight loss preceded death and thus represents a sensitive and quantitative clinical parameter for monitoring disease progression. Comparable patterns have been reported in interferon-deficient mouse models of CCHFV and other viral hemorrhagic fevers, reinforcing the relevance of this endpoint for preclinical evaluation of vaccines and antiviral therapies [6,8,14].

Analysis of viral RNA distribution revealed high viral loads in serum, spleen, and liver, while viral burden in the brain remained comparatively low. This pattern is consistent with earlier studies demonstrating that CCHFV primarily targets peripheral organs involved in immune regulation, hematopoiesis, and metabolism [6,9,17,18]. High viral loads in the spleen likely reflect infection of immune cell populations and may contribute to immune dysregulation and systemic inflammation, while liver involvement is consistent with the hepatic dysfunction commonly observed in severe human CCHF cases [2,9]. The limited viral presence in the brain suggests that neuroinvasion is not a dominant feature of disease in this model, supporting its suitability for studying systemic CCHFV pathogenesis.

The availability of a well-characterized and highly susceptible *in vivo* model is particularly important in the context of the extensive genetic diversity of CCHFV. Multiple phylogenetically distinct genotypes circulate globally, and reassortment events between viral genome segments are frequently observed [4,5,9,16]. Accumulating evidence suggests that genetic variability among CCHFV strains may influence replication efficiency, immune evasion, and virulence; however, experimental comparisons of genovariant-specific pathogenicity remain limited [4,5]. The IFN knockout mouse model described here provides a sensitive system for systematic evaluation of these differences under standardized conditions.

Furthermore, interferon-deficient mouse models have already demonstrated their value in preclinical testing of DNA-based, viral vector, and subunit vaccines, as well as antiviral compounds [8,12–14]. Notably, previous studies have shown that high titers of neutralizing antibodies do not always correlate with protection in IFN-deficient mice, underscoring the importance of robust *in vivo* efficacy testing [13]. The detailed baseline characterization presented in this study establishes a reference framework for future investigations aimed at assessing vaccine-induced protection and antiviral activity against both homologous and heterologous CCHFV genovariants.

Taken together, our findings demonstrate that the combination of SW-13–based virus propagation and IFN knockout mice represents a powerful and versatile experimental platform. This system enables reproducible virus production, reliable quantification, and sensitive assessment of disease outcomes, thereby facilitating advanced studies on CCHFV pathogenic mechanisms and the development of effective medical countermeasures.

## Conclusion

In conclusion, we established an integrated experimental platform for CCHFV research that combines optimized virus propagation in SW-13 cells with a lethal IFN knockout mouse model. Our results demonstrate that CCHFV induces rapid, systemic, and largely dose-independent lethal disease in IFN-deficient mice, accompanied by pronounced weight loss and high viral loads in peripheral organs. The optimized SW-13 cell-based system enables reproducible virus production and reliable titer determination using complementary methods. This well-characterized model provides a robust and versatile tool for future studies aimed at elucidating CCHFV pathogenic mechanisms, comparing the virulence of different viral genovariants, and evaluating vaccines, antiviral compounds, and neutralizing antibodies. Ultimately, such studies will contribute to the development of effective medical countermeasures against Crimean-Congo hemorrhagic fever, a pathogen of increasing global public health concern.

